# Comparative proteomics analysis identifies L-FABP as a putative biomarker of graft injury during liver transplantation

**DOI:** 10.1101/2020.04.22.055442

**Authors:** Maogen Chen, Xiaohong Lin, Yixi Zhang, Qiang Zhao, Peiming Mei, Yinghua Chen, Zhiyong Guo, Linwei Wu, Yi Ma, Dongping Wang, Weiqiang Ju, Xiaoshun He

## Abstract

**Background:** To a large extent the success of liver transplantation depends on quality of allografts. The molecular basis of the susceptibility of different liver allografts to transplant injury remains undefined.

**Methods:** Transplanted liver samples were collected and divided into three groups: the optimal graft (OG) group, early allograft dysfunction (EAD) group, and primary nonfunction (PNF) group. iTRAQ comparative quantitative proteomic analysis and multiple reaction monitoring (MRM) verification was performed.

**Results:** More than 160 differentially expressed proteins were detected in the PNF group, compared to 54 and 36 proteins in the EAD and OG groups respectively. Liver-type fatty acid-binding protein (L-FABP) was found as differentially expressed in both cold preserved and reperfused liver. Serum L-FABP level in donors was higher in the PNF and EAD groups than in the OG group. A lower tissue expression of L-FABP was observed in the PNF groups than other groups after reperfusion, indicating incompetent liver donor quality. In mouse ischemia reperfusion injury (IRI) model, the serum levels and tissue expression of L-FABP corresponded to the ALT variation curve.

**Conclusions:** Suboptimal donor livers are more sensitive to ischemia reperfusion injury. L-FABP might be an effective biomarker for evaluating donor quality in liver transplantation.

## Introduction

Liver transplantation (LT) is a successful and effective technology for the treatment of end-stage liver disease. Due to the scarcity of organ donors, extended criteria donors (ECD) are usually used for patients on waiting lists who require urgent liver transplantation. Marginal liver grafts, such as steatosis allograft and grafts from donation after cardiac death (DCD) donors, usually have a great chance of graft failure and biliary complications, which ultimately increase patient mortality and morbidity. According to differences in postoperative liver function and patient recovery, grafts are divided into three categories: early allograft dysfunction (EAD), primary non-function (PNF) and optimal graft function (OG). Lower quality liver allografts are associated with severe ischemia reperfusion injury (IRI), resulting in the occurrence of EAD or PNF, which significantly affect patient survival and allograft survival (1, 2). The incidence of EAD is up to 39.5% in patients with allografts from DCD donors (1). A high rate of PNF (10%) and retransplantation (15%) in liver transplant patients was reported in a transplant center in Italy (2).

However, until now, there has been a lack of molecular biology methods to evaluate the quality of donor livers. Different liver grafts from donors with high risk factors might induce various degrees of IRI. To identify protein phenotypes associated with liver quality, we performed proteomics analysis of liver IRI in three types of liver graft, namely, the OG, EAD and PNF groups. We further confirmed that a protein, liver-type fatty acid-binding protein (L-FABP), can be used as a liver injury index and might be a marker of donor liver quality.

## Experimental Procedures

### Experimental Design

Thirty-five human liver biopsy samples were used for the iTRAQ and MRM verification methods. Clinical data were collected from 15 liver transplant recipients, and normal liver biopsy samples were acquired from 5 optimal organ donors. All samples were collected from Nov 2015 to Mar 2017 at the First Affiliated Hospital of Sun Yat-sen University (FAH-SYSU), China. Written informed consent was obtained from the recipients. The study protocol conforms to the ethical guidelines of the 1975 Declaration of Helsinki and approved by the Clinical Research (Ethics) Committee of the First Affiliated Hospital of Sun-Yat-Sen University.

The patient cohort was divided into three groups: the PNF, EAD and OG groups. The diagnosis standard of EAD based on serum transaminases levels, total bilirubin levels and the international normalized ratio as previously reported (3, 4). PNF was defined as retransplantation or recipient death resulting from a poor and nonrecoverable allograft function within 7 days after transplantation (5). OG was defined as fast recovery of liver function without rejection and artery or biliary complications.

Liver biopsy samples were obtained at three different time points during liver transplantation. For the normal control group, liver samples were acquired before cold perfusion during the organ harvesting procedure, which was designated as X0. Donor livers were maintained in 4 °C University of Wisconsin solution and transplanted into the recipient after preservation. The second biopsy time point was before donor liver recirculation, which was considered X1. X1 represented the length of cold ischemia time, which resulted in cold preservation injury. After 2 hours of liver artery anastomosis, another liver sample was removed for biopsy at X2. The time period from X0 to X2 indicated the ischemia reperfusion injury phase. All liver tissues were immediately preserved in liquid nitrogen once procured.

### Protein preparation and SDS-PAGE electrophoresis

Protein was extracted from liver tissues with lysis buffer (7 M urea, 2 M thiourea, 4% CHAPS, and 40 mM Tris-HCl, pH 8.5) containing 1 mM PMSF and 2 mM EDTA, and then 10 mM DTT was added 5 min later. These solutions were separated by a tissue lyser and centrifugation at 25,000 × g for 15 min at 4 °C. The supernatant was transferred to a new tube, 10 mM DTT was added to break the disulfide bonds, and the supernatant was incubated in a waterbath at 56 °C for 1 h. Then, 55 mM IAM was added to the solution on ice to block the cysteine residues, and the samples were incubated for 45 min in a darkroom. Chilled acetone was added for sedimentation, and 2 hrs later, the samples were centrifuged at 25,000 × g for 15 min. The protein pellet was dissolved in 0.5 M TEAB and sonicated on ice. After centrifugation at 30,000 × g at 4 °C, the supernatant was collected for testing or preservation at -80 °C for further analysis. The protein concentrations were determined by the Bradford protein quantitation assay.

SDS-PAGE was used to separate proteins and detect their integrity. Thirty micrograms of protein from each sample was loaded on 10% SDS-PAGE gels and Coomassie brilliant blue staining analysis revealed clear band patterns. The samples suitable for subsequent analysis were then subjected to trypsin digestion and LC– MS/MS analysis.

### Quantitative iTRAQ analysis

The protein samples were digested with 2.5 μg trypsin at a protein:trypsin ratio of 40:1 at 37 °C for 4 h, and then trypsin was added for an additional 8 h. The zymolytic peptides were then desalted with a Strata X C18 column and dried by vacuum centrifugation. The peptides were dissolved in 0.5 M TEAB and labeled with 8-plex iTRAQ reagent according to the manufacturer’s protocol. Seven different iTRAQ tags were used to mark the PNF2, PNF1, EAD2, EAD1, OG2, OG1, and control group samples. Briefly, one unit of isobaric tags was dissolved in 24 μL isopropanol. The peptides were incubated with the iTRAQ reagent at room temperature for 2 h and then dried by vacuum centrifugation.

Strong cation exchange (SCX) chromatography was performed with an LC-20AB HPLC Pump system (Shimadzu, Kyoto, Japan). The labeled samples were mixed and reconstituted with 2 mL buffer A (5% ACN, pH 9.8). The peptide mixtures were then loaded onto a 4.6 × 250 mm Gemini C18 column containing 5-μm particles. The peptides were eluted at a flow rate of 1 mL/min with a gradient of buffer B (95% CAN, pH 9.8) for 10 min, 5–35% buffer B for 40 min, and 35–90% buffer B for 1 min. The fractions were collected every minute by measuring the absorbance at 214 nm. Twenty fractions were collected, desalted with a Strata X C18 column and vacuum dried.

Each fraction was dissolved in buffer A (2% ACN, 0.1% FA) and centrifuged at 20,000 × g for 10 min. The supernatant was loaded onto a C18 trap column (inner diameter of 75 μm) in an LC-20 AD nanoHPLC (Shimadzu, Kyoto, Japan) with an autosampler. The samples were loaded at a flow rate of 300 nL/min, and the following gradient procedure was used: 5% buffer B (98% ACN, 0.1% FA) for 8 min followed by a 35-min linear gradient to 35% buffer B, a 5-min linear gradient to 60%, a 2-min linear gradient to 80%, maintenance at 80% B for 5 min, and return to 5% over 1 min. The separation capillary was coupled directly to a mass spectrometer with an electrokinetically driven sheath-flow nanospray interface for further examination.

The fractions were ionized by nano ESI and examined using a tandem mass spectrometer Q-Exactive (Thermo Fisher Scientific, CA, USA) in the DDA (data-dependent acquisition) mode. The nanospray voltage was 1.5 kV. Full MS scans were acquired in the Orbitrap mass analyzer over a 350–1600 m/z range with a mass resolution of 70,000. The 20 most intense multiply charged precursors with molecule-ion intensities greater than 10,000 were selected for higher energy collision-induced dissociation fragmentation. Tandem mass spectra were acquired in the Orbitrap mass analyzer with a mass resolution of 17,500 at 100 m/z. The data were acquired with a 2+ to 7+ charge-state and a 15-s dynamic exclusion setting.

### Data processing and analyses

All primary mass spectrum data were converted to MGF files. The target proteins were identified using the Mascot search engine (Matrix Science, London, UK). Proteins with a 1.2-fold change or greater and p < 0.05 were considered significant when the same result was obtained in more than 3 experiments. Functional annotation of the proteins was performed using the Blast2GO program with comparisons with the nonredundant protein database (NR, NCBI). The KEGG database and the COG (Clusters of Orthologous Groups of Proteins) database were used for protein annotation and enrichment analysis.

### MRM validation of differentially expressed proteins identified by iTRAQ

Sixty proteins identified by ITRAQ were selected for MRM validation. Samples were digested with trypsin as described before and spiked with 50 fmol β-galactosidase for data normalization. MRM analyses were performed on a QTRAP 5500 mass spectrometer (SCIEX, Framingham, MA, USA) equipped with an LC-20AD nanoHPLC system. The mobile phase consisted of solvent A and 0.1% aqueous formic acid and solvent B, 98% acetonitrile and 0.1% formic acid. The peptides were separated on a C18 column (0.075 × 150 mm column, 3.6 μm) at 300 nL/min and eluted with a gradient of 5%–30% solvent B for 38 min, 30%–80% solvent B for 4 min, and maintenance at 80% for 8 min. For the QTRAP5500 mass spectrometer, a spray voltage of 2400 V, a nebulizer gas pressure of 23 p.s.i., and a dwell time of 10 ms were used. Multiple MRM transitions were monitored using unit resolution in both Q1 and Q3 quadrupoles to maximize specificity.

Skyline software was used to integrate the raw file generated by QTRAP 5500 (SCIEX, Framingham, MA, USA). The iRT strategy was adopted to compare a spectrum of a given peptide against a spectral library. All transitions for each peptide were used for quantitation unless interference from the matrix was observed. β-galactosidase was used for label-free data normalization. Using the MS stats with the linear mixed-effects model, the p values were adjusted to maintain the FDR at a cutoff of 0.05. All proteins with a p value below 0.05 and a fold change larger than 1.5 were considered significant.

### Animal management and the model of hepatic ischemia reperfusion injury

Male wild-type C57BL/6J mice (25–30 g, 8–12 weeks of age) were purchased from the Animals Experimentation Center of Sun Yat-sen University (SYSU). The mice were bred in individual cages in an SPF room and were allowed free access to laboratory food and water in sound-attenuated, temperature-controlled isolation chambers at 25 °C ambient temperature. All animal procedures were approved by the Use Committee of Animal Center and ethical committee of FAH-SYSU (Guangzhou, China).

A partial liver ischemia reperfusion injury model was established as described previously (6). The mice were anaesthetized intraperitoneally with pentobarbital sodium. A midline laparotomy was performed to expose the liver. The mice were subjected to occlusion of the hepatic blood supply to the left lateral and median lobe with a microvascular clamp for 30 min followed by 0-24 hrs of reperfusion (except where stated otherwise). After reperfusion the abdominal incision was closed with a 4–0 silk suture. Sham mice were subjected to the same protocol without vascular occlusion. The mice were sacrificed when indicated, and liver tissues and sera specimens were harvested.

### Histopathological and immunohistochemical examination of tissue sections

Mouse liver tissues were fixed in 10% formalin and then embedded in paraffin. The tissues were then cut into slices. Histopathological examination was performed with hematoxylin-eosin staining. The pathological scores were evaluated blindly by two pathologists according to Suzuki’s criteria (7). Immunohistochemistry was also performed to evaluate protein expression using a monoclonal antibody against L-FABP (Abcam, Cambridge, MA). The staining results were captured under a microscope with a camera.

### Analysis of serum L-FABP levels

L-FABP levels in human and mouse sera were measured using enzyme-linked immunosorbent assay kits (R&D Systems Inc., Minneapolis, MN, USA) according to the manufacturer’s recommendations. The minimum detectable level of L-FABP was 0.01 ng/mL.

### Western blot analyses

Protein lysates from liver tissues were homogenized with RIPA lysis buffer and subjected to SDS-PAGE. The proteins were transferred onto polyvinylidene fluoride (PVDF) membranes. After blocking with 5% milk for 1 h, the membranes were stained with a primary antibody against L-FABP overnight at 4 °C followed by a secondary antibody for 1 h at room temperature. The membranes were visualized by an enhanced chemiluminescent kit (Thermo Fisher Scientific).

### Statistical Rationale

All data are presented as the mean±SEM. Statistical analysis was performed with GraphPad Prism 5 software. For statistical analysis of two groups, significant differences were determined by unpaired, two-tailed, Student’s t test. Analysis of variance (ANOVA) was used for analyses of more than two groups. Statistical *p* values <0.05 were considered significant.

## Results

### Demographics and clinical characteristics of recipients and donors

Fifteen liver transplant recipients were chosen for proteomics analysis; they were divided into 3 groups, the PNF, EAD and OG groups, with 5 cases in each group. All patients were male. The median age was 53 (29-67), 54 (34-55) and 53 (46-61) years for the PNF, EAD and OG groups, respectively (Table 1). The mean (± SD) BMI was 20.66 kg/m^2^ (± 2.2) for the PNF group, 26.2 kg/m^2^ (± 1.2) for the EAD group, and 24.2 kg/m^2^ (± 3.7) for the OG group, and 53.3% of recipients had a normal body weight. The major diagnoses of LT patients were liver failure and hepatocellular carcinoma, and there was one case of primary biliary cirrhosis. The preoperative mean MELD scores were 26.8 ± 12.8 for the PNF group, 14.8 ± 7.7 for the EAD group, and 16.6 ± 13.1 for the OG group. The clinical characteristics of LT included cold ischemia time, warm ischemia time, anhepatic time, and surgery and transfusion time and can be found in Table 1. The mean age of the donors was 34.4 ± 7.0 years, 35.8 ± 12.4 years, 22.2 ± 13.9 years for the PNF, EAD and OG groups, respectively. A total of 73.3% of donors were men. The leading primary causes of donor death were traumatic brain injury (60.0%) and adverse cardiovascular events (33.3%) (Table 2).

**Table 1.**
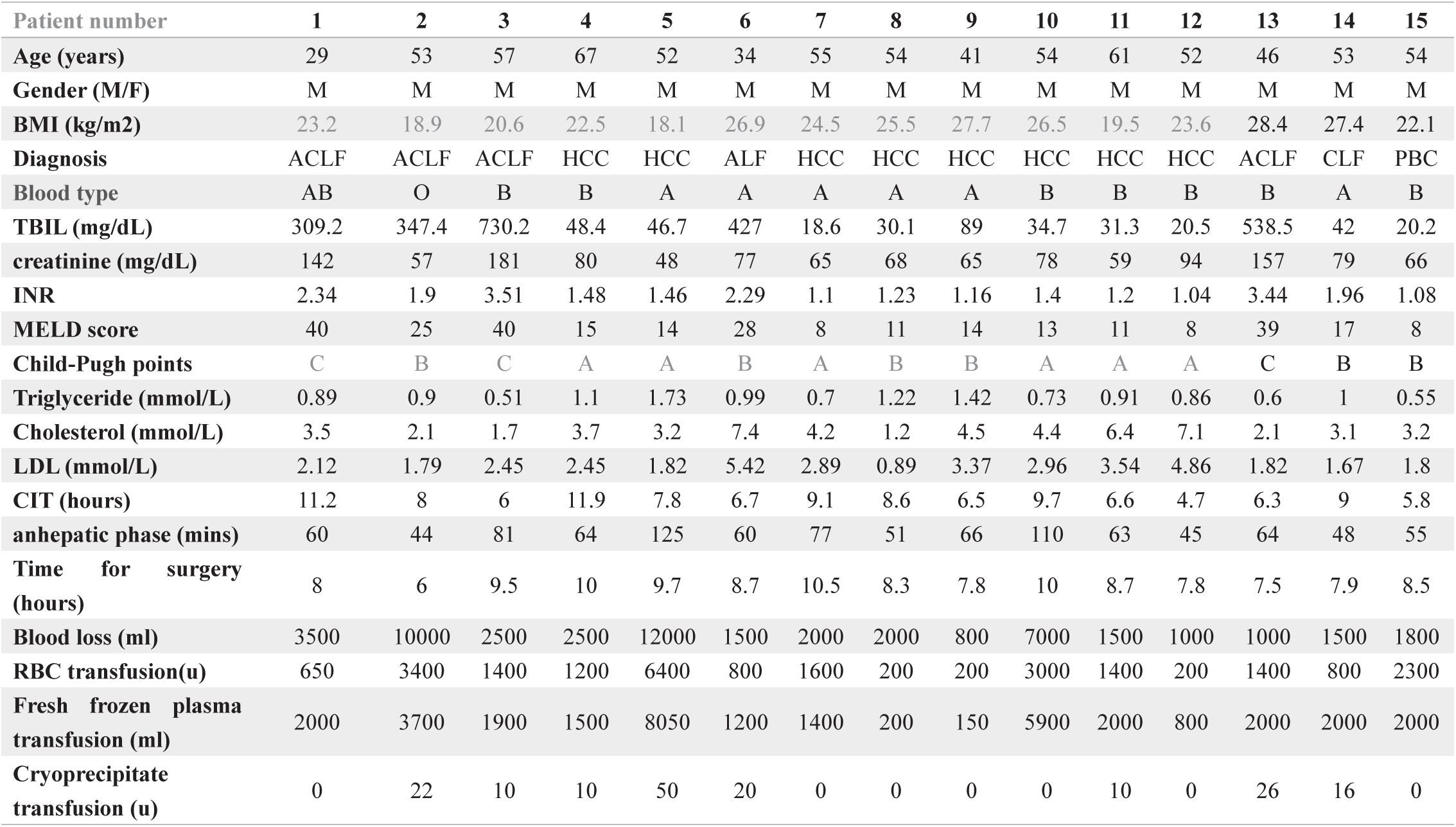
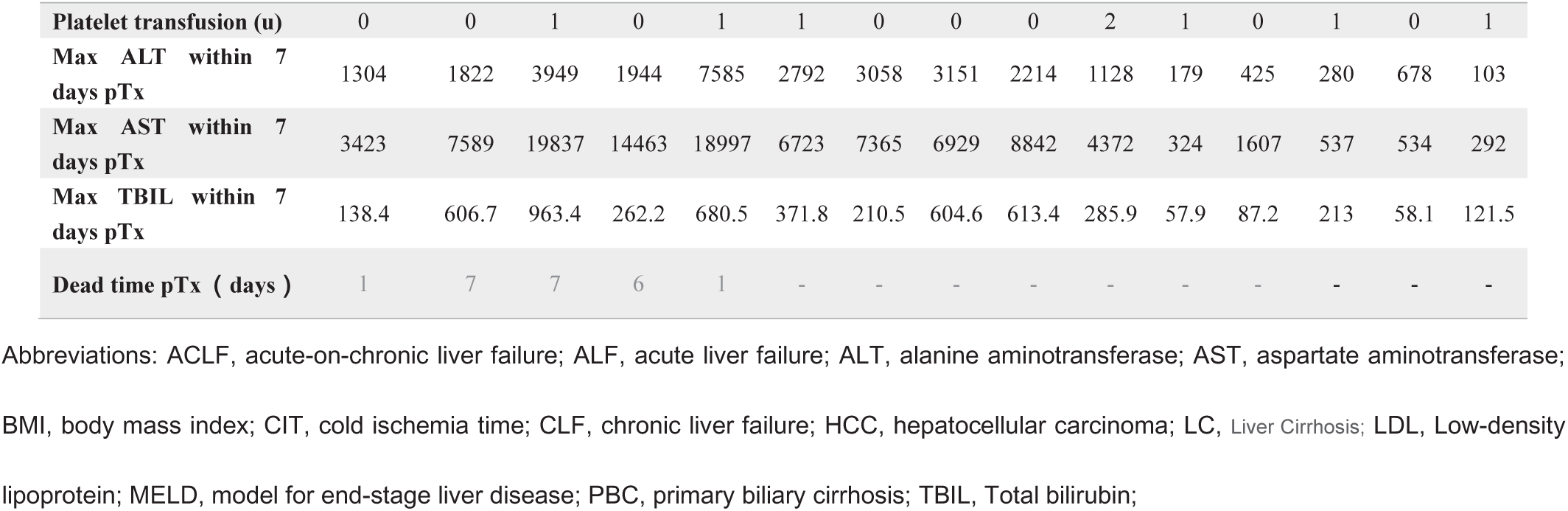
Recipients demographic data and clinical Characteristics.

| Patient number | 1 | 2 | 3 | 4 | 5 | 6 | 7 | 8 | 9 | 10 | 11 | 12 | 13 | 14 | 15 |
| --- | --- | --- | --- | --- | --- | --- | --- | --- | --- | --- | --- | --- | --- | --- | --- |
| Age (years) | 29 | 53 | 57 | 67 | 52 | 34 | 55 | 54 | 41 | 54 | 61 | 52 | 46 | 53 | 54 |
| Gender (M/F) | M | M | M | M | M | M | M | M | M | M | M | M | M | M | M |
| BMI (kg/m2) | 23.2 | 18.9 | 20.6 | 22.5 | 18.1 | 26.9 | 24.5 | 25.5 | 27.7 | 26.5 | 19.5 | 23.6 | 28.4 | 27.4 | 22.1 |
| Diagnosis | ACLF | ACLF | ACLF | HCC | HCC | ALF | HCC | HCC | HCC | HCC | HCC | HCC | ACLF | CLF | PBC |
| Blood type | AB | O | B | B | A | A | A | A | A | B | B | B | B | A | B |
| TBIL (mg/dL) | 309.2 | 347.4 | 730.2 | 48.4 | 46.7 | 427 | 18.6 | 30.1 | 89 | 34.7 | 31.3 | 20.5 | 538.5 | 42 | 20.2 |
| creatinine (mg/dL) | 142 | 57 | 181 | 80 | 48 | 77 | 65 | 68 | 65 | 78 | 59 | 94 | 157 | 79 | 66 |
| INR | 2.34 | 1.9 | 3.51 | 1.48 | 1.46 | 2.29 | 1.1 | 1.23 | 1.16 | 1.4 | 1.2 | 1.04 | 3.44 | 1.96 | 1.08 |
| MELD score | 40 | 25 | 40 | 15 | 14 | 28 | 8 | 11 | 14 | 13 | 11 | 8 | 39 | 17 | 8 |
| Child-Pugh points | C | B | C | A | A | B | A | B | B | A | A | A | C | B | B |
| Triglyceride (mmol/L) | 0.89 | 0.9 | 0.51 | 1.1 | 1.73 | 0.99 | 0.7 | 1.22 | 1.42 | 0.73 | 0.91 | 0.86 | 0.6 | 1 | 0.55 |
| Cholesterol (mmol/L) | 3.5 | 2.1 | 1.7 | 3.7 | 3.2 | 7.4 | 4.2 | 1.2 | 4.5 | 4.4 | 6.4 | 7.1 | 2.1 | 3.1 | 3.2 |
| LDL (mmol/L) | 2.12 | 1.79 | 2.45 | 2.45 | 1.82 | 5.42 | 2.89 | 0.89 | 3.37 | 2.96 | 3.54 | 4.86 | 1.82 | 1.67 | 1.8 |
| CIT (hours) | 11.2 | 8 | 6 | 11.9 | 7.8 | 6.7 | 9.1 | 8.6 | 6.5 | 9.7 | 6.6 | 4.7 | 6.3 | 9 | 5.8 |
| anhepatic phase (mins) | 60 | 44 | 81 | 64 | 125 | 60 | 77 | 51 | 66 | 110 | 63 | 45 | 64 | 48 | 55 |
| Time for surgery (hours) | 8 | 6 | 9.5 | 10 | 9.7 | 8.7 | 10.5 | 8.3 | 7.8 | 10 | 8.7 | 7.8 | 7.5 | 7.9 | 8.5 |
| Blood loss (ml) | 3500 | 10000 | 2500 | 2500 | 12000 | 1500 | 2000 | 2000 | 800 | 7000 | 1500 | 1000 | 1000 | 1500 | 1800 |
| RBC transfusion(u) | 650 | 3400 | 1400 | 1200 | 6400 | 800 | 1600 | 200 | 200 | 3000 | 1400 | 200 | 1400 | 800 | 2300 |
| Fresh frozen plasma transfusion (ml) | 2000 | 3700 | 1900 | 1500 | 8050 | 1200 | 1400 | 200 | 150 | 5900 | 2000 | 800 | 2000 | 2000 | 2000 |
| Cryoprecipitate transfusion (u) | 0 | 22 | 10 | 10 | 50 | 20 | 0 | 0 | 0 | 0 | 10 | 0 | 26 | 16 | 0 |
| **Platelet transfusion (u)** | 0 | 0 | 1 | 0 | 1 | 1 | 0 | 0 | 0 | 2 | 1 | 0 | 1 | 0 | 1 |
| **Max ALT within 7 days pTx** | 1304 | 1822 | 3949 | 1944 | 7585 | 2792 | 3058 | 3151 | 2214 | 1128 | 179 | 425 | 280 | 678 | 103 |
| **Max AST within 7 days pTx** | 3423 | 7589 | 19837 | 14463 | 18997 | 6723 | 7365 | 6929 | 8842 | 4372 | 324 | 1607 | 537 | 534 | 292 |
| **Max TBIL within 7 days pTx** | 138.4 | 606.7 | 963.4 | 262.2 | 680.5 | 371.8 | 210.5 | 604.6 | 613.4 | 285.9 | 57.9 | 87.2 | 213 | 58.1 | 121.5 |
| **Dead time pTx ( days )** | 1 | 7 | 7 | 6 | 1 | - | - | - | - | - | - | - | - | - | - |

Abbreviations: ACLF, acute-on-chronic liver failure; ALF, acute liver failure; ALT, alanine aminotransferase; AST, aspartate aminotransferase; BMI, body mass index; CIT, cold ischemia time; CLF, chronic liver failure; HCC, hepatocellular carcinoma; LC, Liver Cirrhosis; LDL, Low-density lipoprotein; MELD, model for end-stage liver disease; PBC, primary biliary cirrhosis; TBIL, Total bilirubin;

**Table 2.**
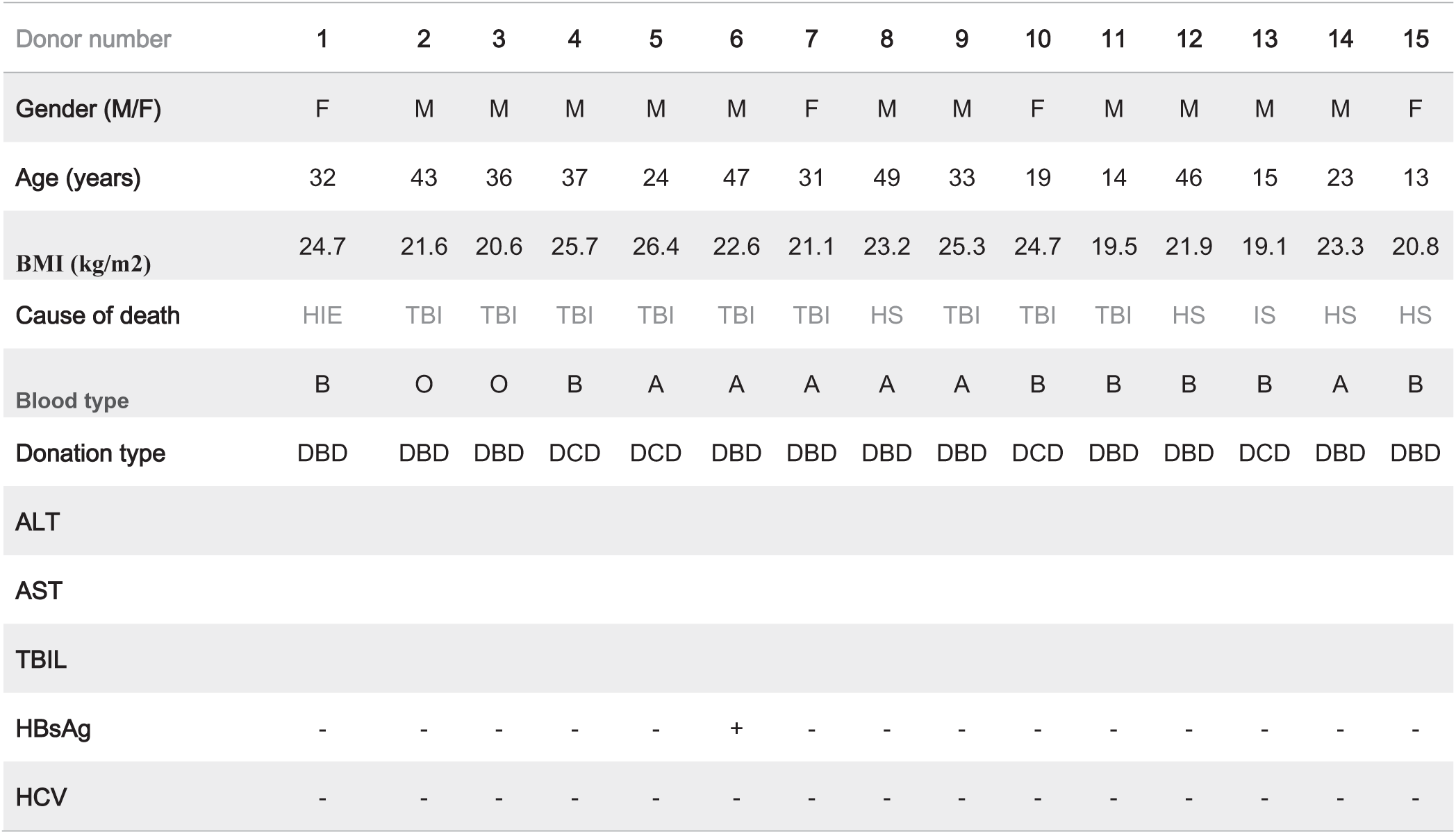
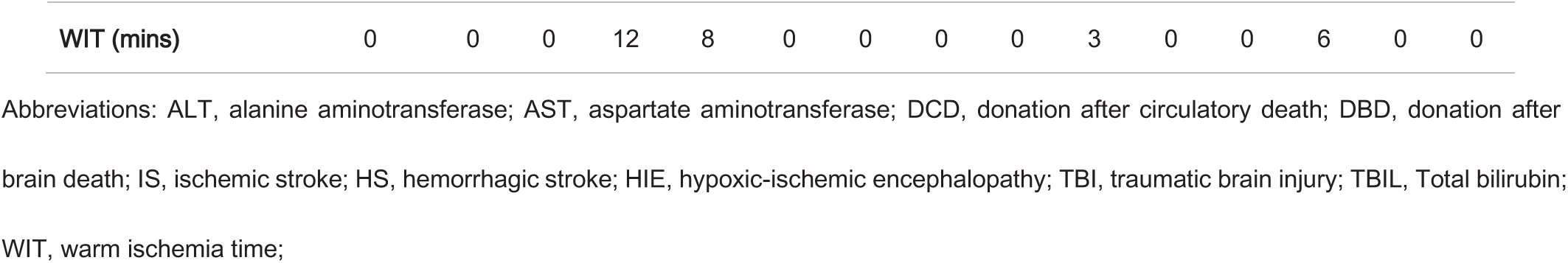
Clinical Features of Organ Donors.

| Donor number | 1 | 2 | 3 | 4 | 5 | 6 | 7 | 8 | 9 | 10 | 11 | 12 | 13 | 14 | 15 |
| --- | --- | --- | --- | --- | --- | --- | --- | --- | --- | --- | --- | --- | --- | --- | --- |
| Gender (M/F) | F | M | M | M | M | M | F | M | M | F | M | M | M | M | F |
| Age (years) | 32 | 43 | 36 | 37 | 24 | 47 | 31 | 49 | 33 | 19 | 14 | 46 | 15 | 23 | 13 |
| BMI (kg/m<sup>2</sup>) | 24.7 | 21.6 | 20.6 | 25.7 | 26.4 | 22.6 | 21.1 | 23.2 | 25.3 | 24.7 | 19.5 | 21.9 | 19.1 | 23.3 | 20.8 |
| Cause of death | HIE | TBI | TBI | TBI | TBI | TBI | TBI | HS | TBI | TBI | TBI | HS | IS | HS | HS |
| Blood type | B | O | O | B | A | A | A | A | A | B | B | B | B | A | B |
| Donation type | DBD | DBD | DBD | DCD | DCD | DBD | DBD | DBD | DBD | DCD | DBD | DBD | DCD | DBD | DBD |
| ALT |  |  |  |  |  |  |  |  |  |  |  |  |  |  |  |
| AST |  |  |  |  |  |  |  |  |  |  |  |  |  |  |  |
| TBIL |  |  |  |  |  |  |  |  |  |  |  |  |  |  |  |
| HBsAg | - | - | - | - | - | + | - | - | - | - | - | - | - | - | - |
| HCV | - | - | - | - | - | - | - | - | - | - | - | - | - | - | - |
| **WIT (mins)** | 0 | 0 | 0 | 12 | 8 | 0 | 0 | 0 | 0 | 3 | 0 | 0 | 6 | 0 | 0 |

Abbreviations: ALT, alanine aminotransferase; AST, aspartate aminotransferase; DCD, donation after circulatory death; DBD, donation after brain death; IS, ischemic stroke; HS, hemorrhagic stroke; HIE, hypoxic-ischemic encephalopathy; TBI, traumatic brain injury; TBIL, Total bilirubin; WIT, warm ischemia time;

### Proteome of the human liver during transplantation

In the present study, quantitative proteomic analysis (iTRAQ) was used to identify proteins that accumulated at different time points in donor livers before and after reperfusion during liver transplantation. The proteomics analysis flow-chart is shown in supplemental Fig. S1. A total of 1,719,726 spectra were obtained, of which 41,178 unique peptides and 6,505 proteins were matched to known spectra in the reference genomes using Mascot software.

A total of 6,048 proteins were functionally annotated with Gene Ontology (GO) terms. 5,402 of the proteins participate in biological processes with 33 unique proteins, most of which are associated with the GO terms single-organism processes, metabolic processes, cellular processes and biological regulation. A total of 1,256 proteins engaged in molecular functions (including 91 unique proteins) were identified, with the most proteins being associated with the GO terms catalytic activity and binding processes. The number of cellular components and unique cellular components represented by the proteins was 5,782 and 361, respectively, with the most proteins being associated with the GO terms cell, cell part, organelle, organelle part, and membrane (Fig. 1A). 4,886 proteins were associated with all three GO terms (Fig. 1B).

**Fig. 1.**
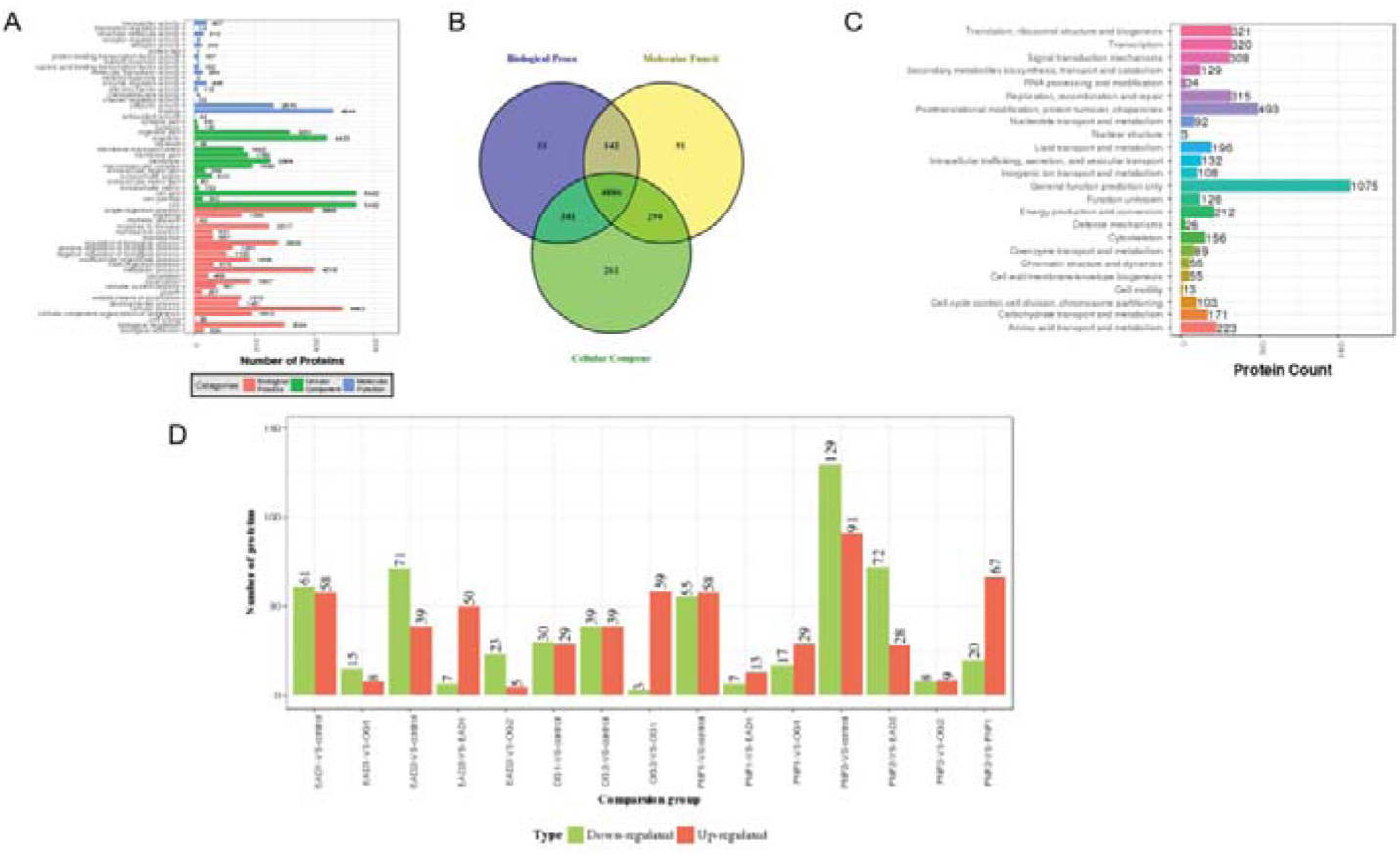
Differential proteins were analyzed by proteomics data in human liver tissues during transplantation. Proteins were functionally annotated with Gene Ontology (GO) for biological process, molecular function, and cellular component (A), and a Venn diagram of the annotation of the proteins for the three processes was created (B). Proteins were divided by COG functional classification (C). Differentially expressed proteins between the groups are shown (D). Red column: upregulated proteins; Green column: downregulated proteins.

A total of 5,977 proteins (98.83%) were annotated by KEGG pathway analysis, and 324 pathways were identified (Fig. 1C). Many proteins were found to participate in the processes of metabolic pathways (15.11%), endocytosis (3.6%), the PI3K-Akt signaling pathway (3.05%), phagosomes (2.64%), protein processing in the endoplasmic reticulum (2.43%), spliceosomes (2.28%), regulation of the actin cytoskeleton (2.24%), focal adhesion (2.21%) and RNA transport (2.16%). There were 182, 107, 98, 84 and 82 proteins identified in the PI3K-Akt signaling pathway, the MAPK signaling pathway, the Ras signaling pathway, the mTOR signaling pathway and apoptosis pathways, respectively. After comparative proteomics analysis of pairs among the 7 groups (the PNF1, PNF2, EAD1, EAD2, OG1, OG2, and control groups), 579 differentiated proteins were found to be upregulated or downregulated (Fig. 1D).

### Differentially expressed proteins after liver ischemia reperfusion injury

Specific proteins associated with IRI were identified by comparing the PNF2, EAD2, OG2 and control groups. Comparisons of iTRAQ proteomics results of the PNF2 and control groups, EAD2 and control groups and OG2 and control groups identified a total of 220, 110 and 78 differentially expressed proteins, including 91 upregulated and 129 downregulated proteins, 39 upregulated and 71 downregulated proteins, 39 upregulated and 39 downregulated proteins, respectively (Fig. 2A). Twenty-two proteins were differentially expressed in the three pairs, including RIDA, F10A1, GSTO1, KCY, MDHC, PARK7, AK1A1, FKB1A, PSB7, L-FABP, HINT1, TRFL, CX7A2, GCSH, PERM, ANXA3, HRG, NGAL, ARF6, LYSC, MMP9, and PRTN3 (Fig. 2B). Based on annotation with GO terms, 21 proteins are involved in biological processes, including cellular processes, metabolic processes, single-organism processes, biological regulation, response to stimulus, cellular component organization or biogenesis, localization, and multicellular organismal processes (Fig. 2C). All 22 proteins plays a role in cellular components and molecular function and most are represented by the terms cell part, organelle part for cellular component and binding, and catalytic activity for molecular function. These proteins are involved in the following pathways: the metabolic pathway, the IL-17 signaling pathway, fat digestion and absorption, the Ras signaling pathway, the TNF signaling pathway, the PPAR signaling pathway, the estrogen signaling pathway, et al.

**Fig. 2.**
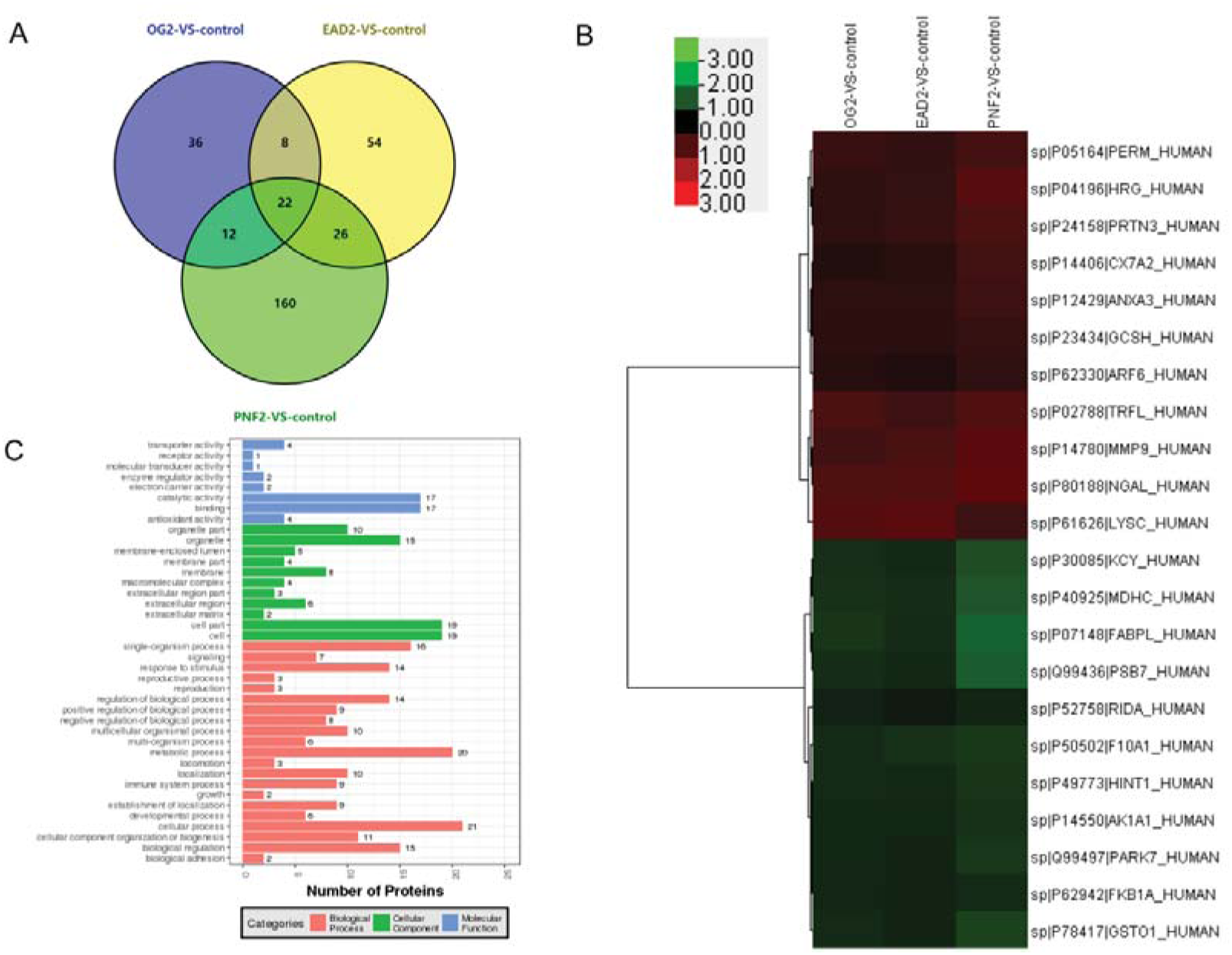
Comparison of proteins identified in the control and tissue groups after reperfusion, which might play an important role in IRI pathogenesis. A shows a Venn diagram of differentially expressed proteins between PNF2, EAD2, OG2 and control liver tissue. B shows cluster analysis of the common proteins that were differentially expressed in the three groups. C represents the GO annotation analysis results of the differentially expressed proteins.

### Protein annotation after cold preservation of donor livers

Differential proteins related to cold preservation damage were also analyzed by comparing the PNF1, EAD1, and OG1 groups to the control group. A total of 202 proteins were found to be differentially expressed among those groups, 17 of which were differentially expressed between all of the groups (Fig. 3A). The differentially expressed proteins were as follows: F10A1, BLVRB, GDIR2, ZN648, HBB, ZA2G, HBA, PRDX2, L-FABP, SC23A, ACS2A, GBB2, THIK, TBA4A, RL12, SEC13, and BDH (Fig. 3B). All 17 proteins are involved in biological processes, the cellular component and molecular function based on GO term annotation analysis. Most of the representative proteins participate in binding and catalytic activity for molecular function, cell and cell part for cellular component, and metabolic processes and cellular processes for biological processes (Fig. 3C). For pathway analysis, the following pathways were found to be involved in the process of cold preservation injury: the PPAR signaling pathway, the mTOR signaling pathway, the Ras signaling pathway, apoptosis, chemokine signaling pathways, the PI3K-Akt signaling pathway, et al.

**Fig. 3.**
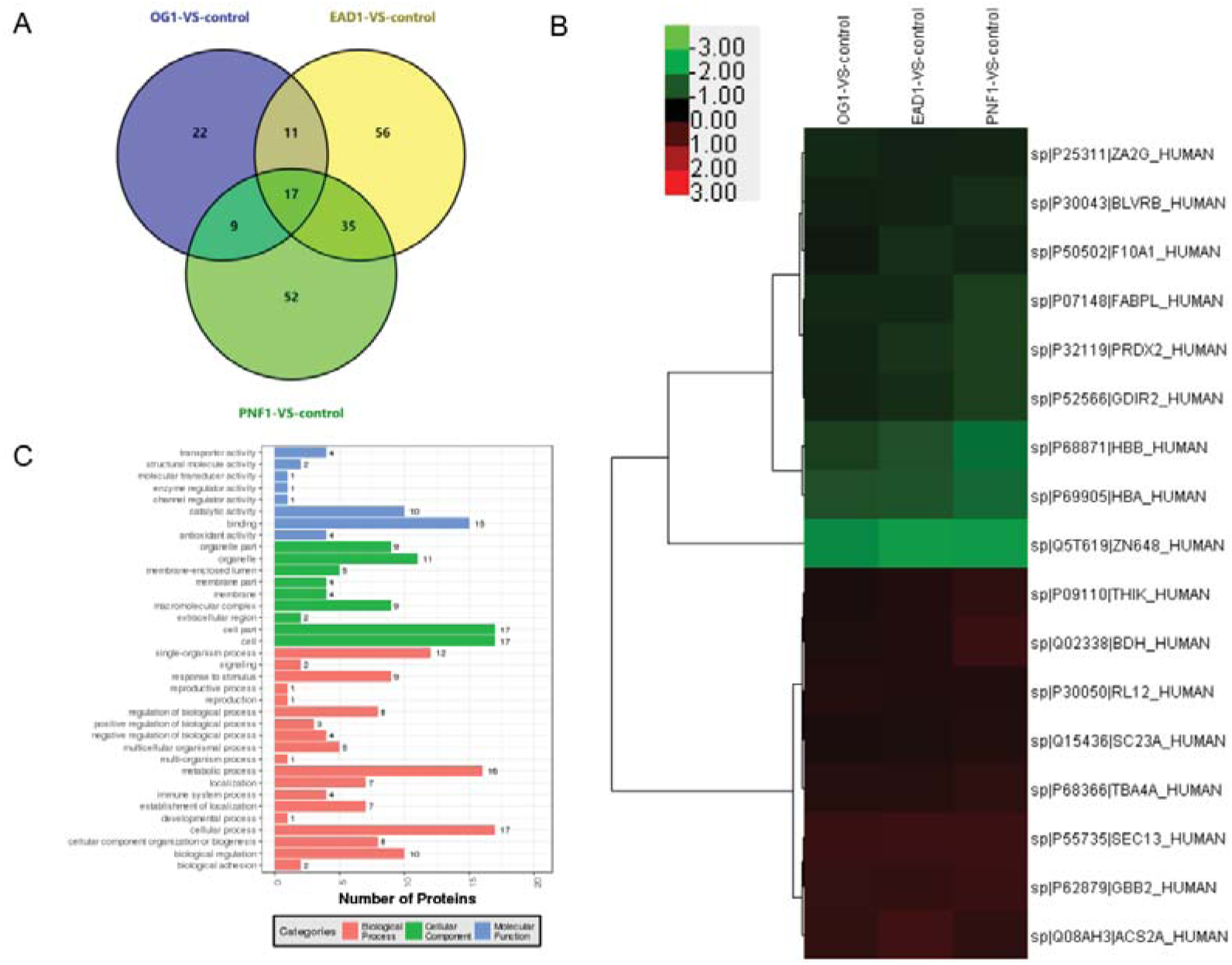
Comparative proteins were identified in the control and cold preservation groups and might represent markers of cold storage injury. A shows a Venn diagram of differentially expressed proteins between PNF1, EAD1, OG1 and control liver tissue. B shows cluster analysis of the common proteins that were differentially expressed in all three groups. C represents the GO annotation analysis results of the differentially expressed proteins.

### MRM validation of differentially expressed proteins

MRM analysis was used to validate the target proteins expressed in the liver. In the present study, 119 proteins were chosen for MRM verification, and 103 proteins were suitable for MRM analysis. Finally, 57 proteins with a high level of fold change and association with liver injury were used for MRM verification. The log ratios of the proteins determined by MRM quantitative analysis were significantly positively correlated with the ratios determined by iTRAQ (Fig. 4A, R = 0.6465). In the comparison between the PNF2 group and the control group, the significant upregulation of 1 protein (VWF) and the significant downregulation of 8 proteins (L-FABP, THIO, HPPD, 6PGD, ACY1, PRDX1, HGD, and CNDP2) identified by iTRAQ were consistent with the results of MRM analysis.

**Fig. 4.**
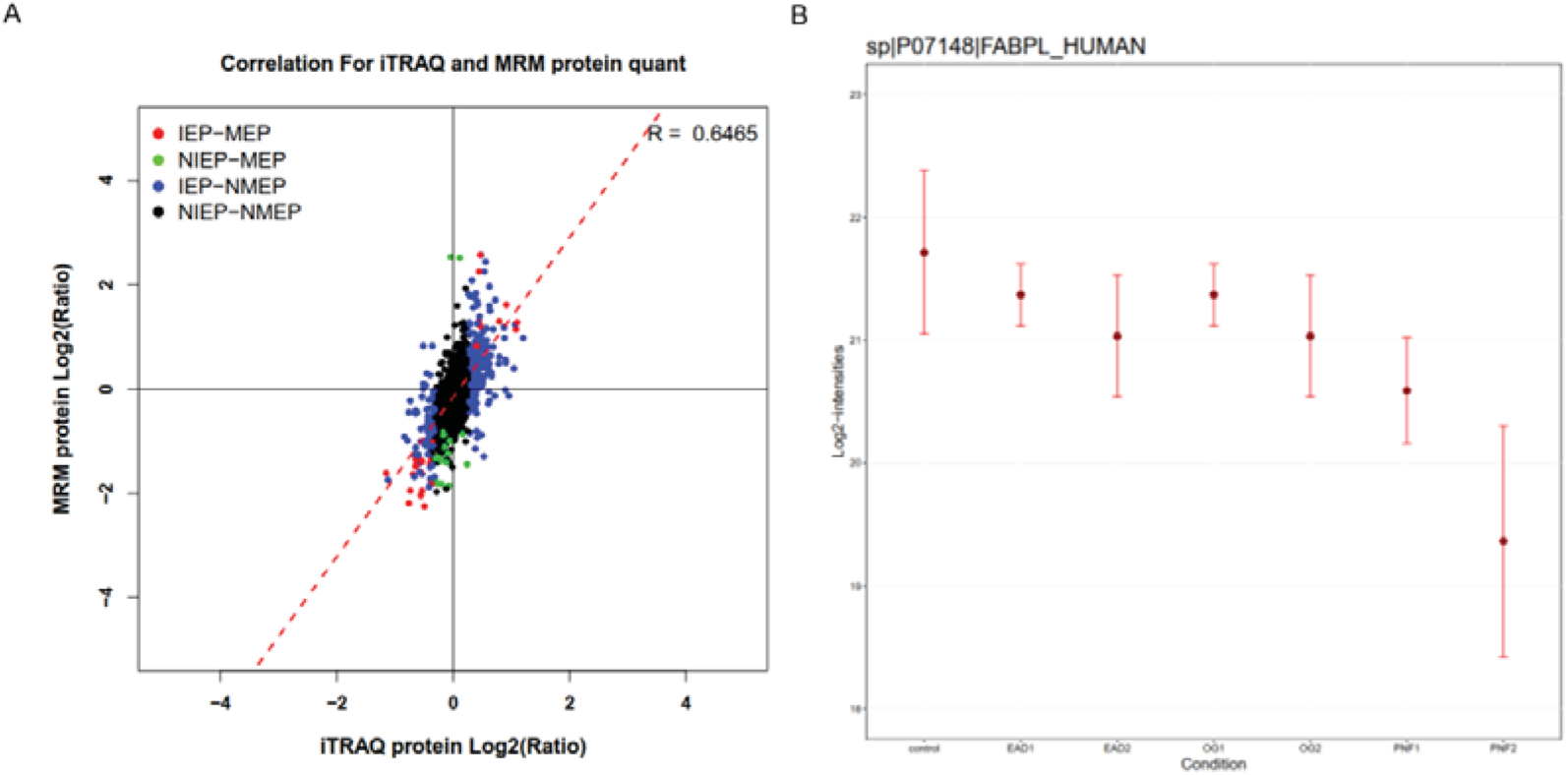
L-FABP might be a possible marker to identify liver donor with lower quality. A. The correlation between iTRAQ and MRM quantified log ratio for target proteins of the comparison groups. B. Normalized peptide intensity of FABPL in all groups of MRM confirmation analysis.

### L-FABP is a differential protein in human transplantation

We further compared proteins between the X1 and control groups and the X2 and control groups, evaluating cold preservation injury- and IRI-related proteins separately. The results showed that 2 proteins, L-FABP and F10A1, were significantly differentially expressed in injured liver tissue and normal liver tissue (supplemental Fig. S2). In the subsequent MRM validation process, the protein intensity of L-FABP in all groups showed that its expression in the PNF1 group was significantly lower than that in the EAD1, OG1 and control groups. L-FABP expression in the PNF2 group was the lowest among all groups (Fig. 4B). As shown above, L-FABP is a differentially expressed protein in both cold preservation injury and reperfusion injury, indicating that it is a possible marker of liver quality.

### L-FABP expression is increased after IRI injury in the mouse

Since differences of L-FABP expression existed before and after reperfusion, it might play a role during the pathological process in IRI. An IRI mouse model was used to confirm this result. After 30 min of 70% hepatic I/R in mice, the levels of ALT and AST were significantly increased, and they reached the highest levels after 6 hours of reperfusion (Fig. 5A). Regarding pathological evaluation, the sham operation group exhibited normal liver structure, whereas IRI caused obvious necrosis, hemorrhage and inflammation infiltration in the liver, as shown by the Suzuki scores (Fig. 5B, C).

**Fig. 5.**
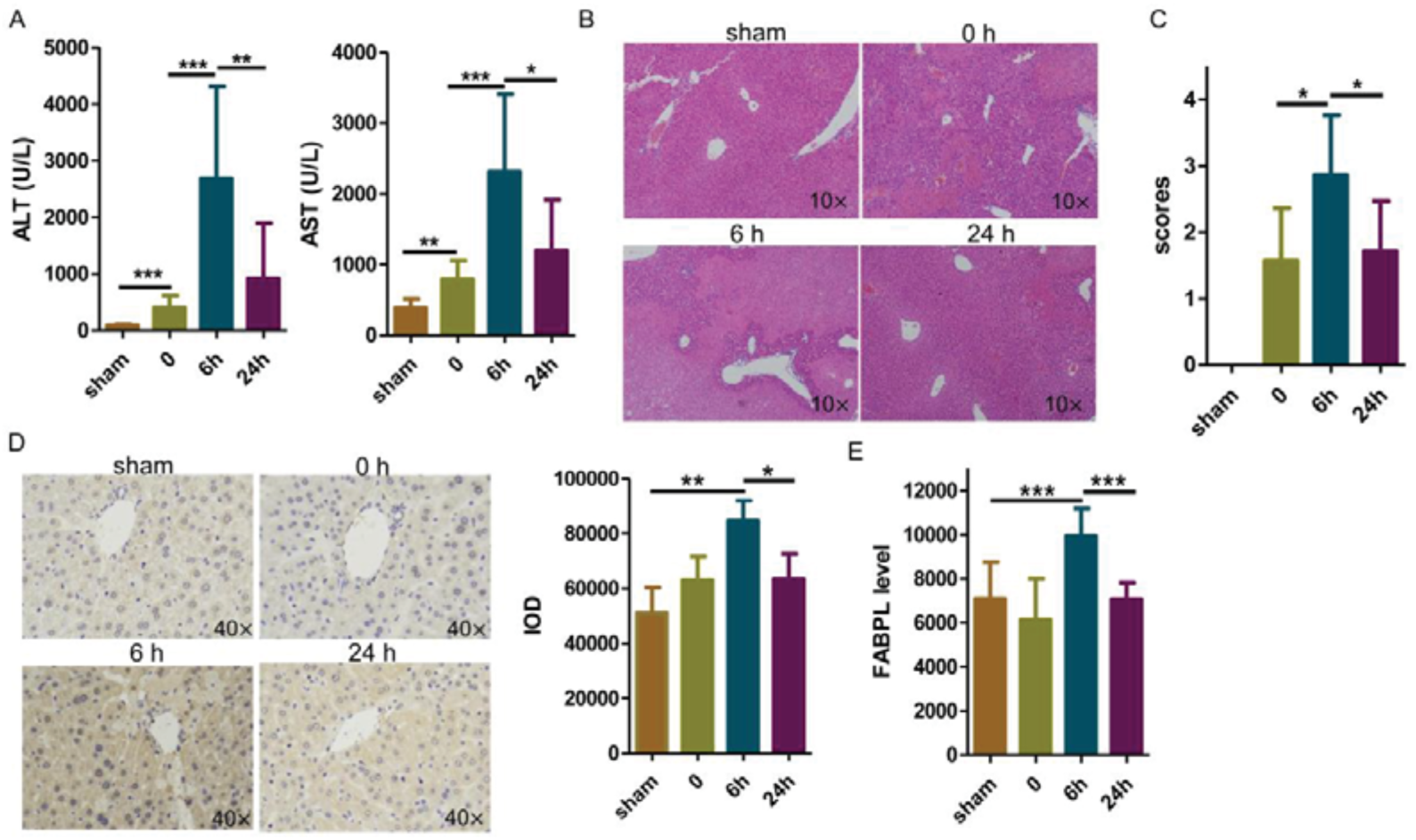
L-FABP is a biomarker of liver injury in a mouse IRI model. (A) Serum ALT and AST levels were determined after different reperfusion times in the IRI group and sham group. (B) HE staining of liver sections from the IRI groups at different time points. Original magnification: 100×. (C) The mean pathological score of the liver sections. All values are expressed as the mean ± SEM. n = 6 independent sections. **P<0.05. (D) Immunohistochemical staining of liver specimens with an antibody against mouse L-FABP is shown. The mean density of positive staining was calculated, as shown in the right panel. (E) Serum L-FABP levels in IRI mice were examined by an ELISA kit. The data are presented as the mean ± SEM. *P<0.05, **P<0.01, and ***P<0.001 between two groups, as indicated.

The time point at which the most severe injury was observed was 6 hours after reperfusion, which is consistent with the change in liver function markers. Immunohistochemistry indicated that L-FABP expression was significantly induced by 6 h of I/R in mice (Fig. 5D). The level of L-FABP in the mouse serum was further examined, and the results showed that L-FABP increased after 6 hours of reperfusion but decreased back to normal levels after 24 hours (Fig. 5E). This result indicates that, like ALT and AST, L-FABP is a marker of liver injury in mice.

### L-FABP might be a sign of liver donor quality in humans

To investigate the function of L-FABP in human liver transplantation, we first checked the L-FABP level in the serum of LT donors and recipients. Since marginal donor factors would affect donor liver recovery, donor sera were tested for L-FABP levels before procurement, and the results showed that the donor L-FABP levels in both PNF and EAD group were significantly higher than that in the OG group (Fig. 6A). Unfortunately, there was no difference in donor L-FABP levels between the PNF group and the EAD group. After reperfusion, the L-FABP level was also higher in both the PNF and EAD groups than in the OG group (Fig. 6B), suggesting that it is a marker of liver injury. Histopathology of liver donors showed higher expression of L-FABP after reperfusion injury in the OG and EAD groups compared to the normal PNF group (Fig. 6C, D). However, L-FABP expression in hepatocytes after cold preservation was not different between those groups. In a word, higher serum level and lower tissue L-FABP expression was a suboptimal marker of donor liver quality.

**Fig. 6.**
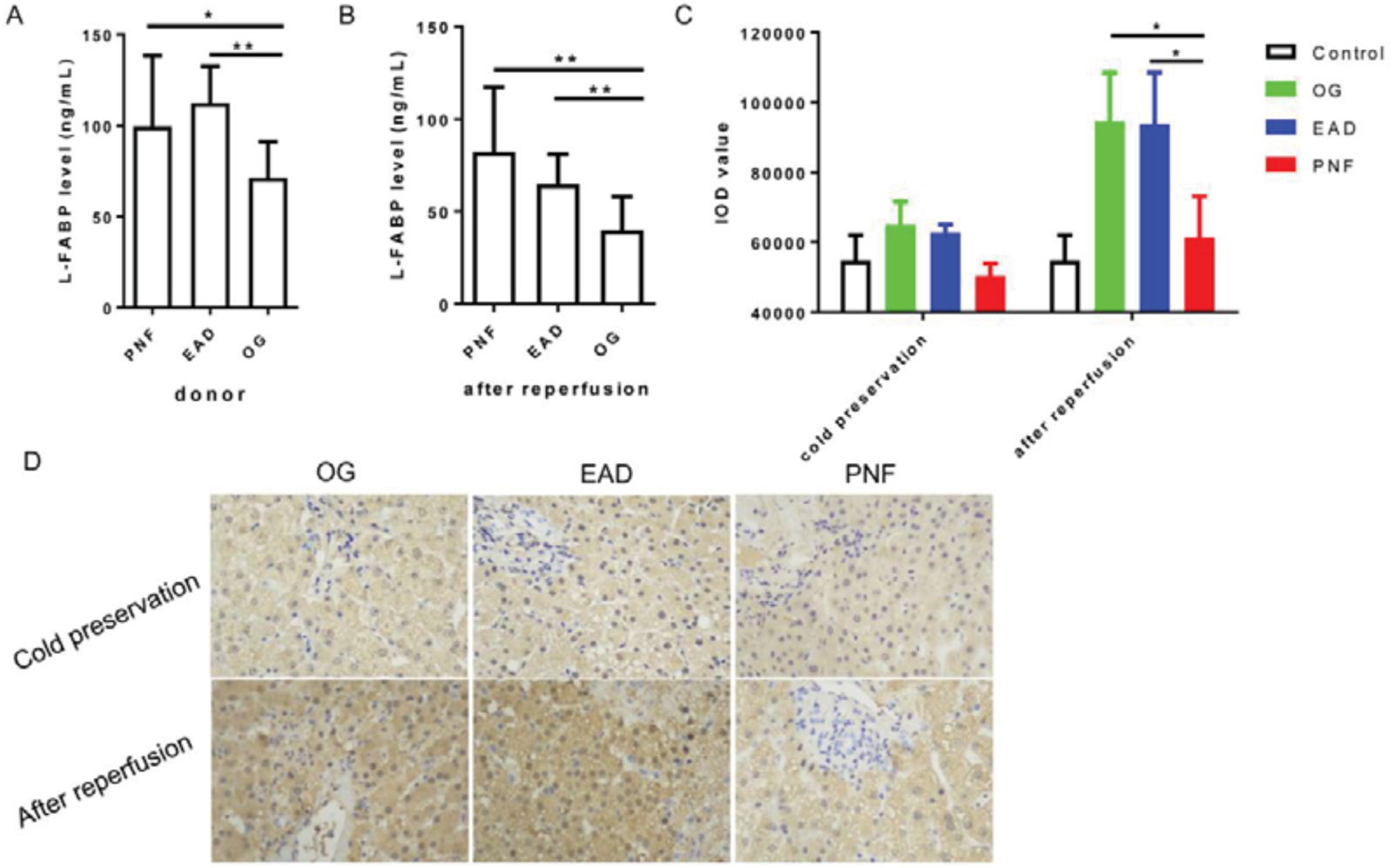
L-FABP might be a suboptimal donor liver injury index for human liver transplantation. (A) Liver transplant recipients were selected for serum L-FABP detection during the first week of surgery. (B) Donor sera acquired before organ procurement were investigated for L-FABP concentration, and differences among the PNF, EAD and OG groups were compared. (C) After 2 hours of donor liver reperfusion, recipient serum L-FABP levels were examined by ELISA. (D, E) L-FABP expression in liver tissue after cold preservation and reperfusion and in normal control tissue was evaluated by immunohistochemistry. Panel D shows representative data of L-FABP expression in hepatocyte cytoplasm. Panel E shows the relative expression of L-FABP in tissues from the different groups. The data are presented as the mean ± SEM. ns, no significance; \**P*<0.05, \*\**P*<0.01, and ***P<0.001 between two groups, as indicated.

## Discussion

To overcome the high incidence of EAD or PNF, an accurate assessment of graft function is required. At present, marginal or ECD liver grafts are considered to be at high risk for graft dysfunction, biliary complications, relapse of hepatitis, and even graft loss or patient death (8). According to the European Liver Transplant Association, suboptimal ECD donors are characterized by the following parameters: donor age ≥65, ICU stay with ventilation ≥7 days, BMI ≥30 kg/m^2^, liver steatosis ≥40%, serum sodium levels ≥165 mmol/L, ALT levels ≥105 U/L, AST levels ≥ 90U/L), and bilirubin levels ≥3 mg/dl) (9). Currently, approximately 8% of liver graft candidates on the waiting list die each year, and 22.3% of recipients have to accept ECD livers (8). High mortality (35.3%) within the first month is observed in recipients with high MELD scores that received ECD grafts, and the six months postoperative graft survival rate using marginal grafts is was 78.9%, whereas it is 93.22% for optimal grafts (8, 10). Improving the quality of ECD liver grafts is an important research direction. Despite the uncertainty of their accuracy, the donor risk index and extended-criteria donor scores have been reported to be helpful in assessing liver graft quality (8). When the quality of a liver graft is questionable, hypothermic oxygenated machine perfusion (HOPE) and normothermic machine perfusion (NMP) are feasible procedures for improving graft quality (11-13). In brief, good transplant recovery depends on accurate liver donor quality evaluation before surgery.

Finding some molecular markers would help to identify suboptimal donor livers for transplant. Our proteomics data showed that 17 proteins, F10A1, BLVRB, GDIR2, ZN648, HBB, ZA2G, HBA, PRDX2, L-FABP, SC23A, ACS2A, GBB2, THIK, TBA4A, RL12, SEC13, and BDH, are differentially expressed after cold preservation, and that these proteins may be markers for quality evaluation. However, some other papers have also reported related results due to different model or injury. Xie H, et al (14) has reported that several proteins, including NOX-1, ICAM-1, cytochrome C, and VEGF, are an independent EAD predictor, since they expressed significantly higher in hepatocytes in pretransplant biopsy specimens via immunohistochemical analysis. Quantitative proteome analysis showed that, in the process of acetaminophen overdose-induced acute liver injury, decreased proteins Prdx6, Prdx3, Aldh2, Gzmf, Bax, Apaf-1, stat3, P4hα1, Ncam, α-SMA, and Cygb participate in the process of liver injury or repair (15). It still need more investigations for confirmation of the liver quality markers.

Hepatocytes can secrete some small proteins in response to liver damage including liver-type fatty acid-binding protein (L-FABP; 14 kDa), α-glutathione S-transferase (α-GST; 26 kDa), alanine transaminase (ALT; 96 kDa) and aspartate transaminase (16). Circulating L-FABP levels increase earlier than α-GST and ALT levels during acute rejection of liver transplantation (16), indicating that L-FABP a sensitive biomarker of liver injury. L-FABP regulates intracellular fatty acid homeostasis via binding and transporting them to mitochondria or peroxisomes for β-oxidization (17). The overexpression of L-FABP in liver cells reduces oxidative stress injury in vitro, suggesting it is a strong endogenous antioxidant (18). The upstream regulatory genes of L-FABP include hepatocyte retinoid X receptor α (RXRα) (19) and ischemia-oxidative stress-related genes such as like hypoxia-inducible factor-1 (HIF-1) and hepatocyte nuclear factor (HNF-1, HNF-4) (20, 21). L-FABP deletion in liver tissue protects against high saturated fat induced steatosis and fibrosis (22). Our data confirmed that L-FABP is an injury marker during human liver transplantation and in a mouse IRI model.

The plasma level of L-FABP is significantly elevated in drug-induced and alcohol-induced liver injury, liver carcinoma and intestinal ischemia injury (23). L-FABP has been proposed to be a sensitive biomarker of hepatocellular damage during drug-induced liver injury, chronic hepatitis C infection and liver transplantation (16, 24, 25). In liver resection surgery with intermittent Pringle maneuver, L-FABP is released from hepatocytes and the circulation levels increase after the completion of liver transection and decrease rapidly (26). Although L-FABP abundantly exists in intestinal villi, it seems that short-term portal vein occlusion during the anhepatic phase of liver resection or transplantation would not result in intestine-derived L-FABP production (27). Considering its low molecular mass and short plasma half-life of 11 min, L-FABP may be a complementary biomarker for liver injury to the conventional marker of ALT levels (24, 25). Our data showed that a higher level of serum L-FABP before donation or after reperfusion might be an effective index for the occurrence of PNF or EAD after liver transplantation. Additionally, lower expression of L-FABP in liver tissue after liver reperfusion might increase the possibility of PNF occurrence after transplantation. So, serum L-FABP implies a marker of suboptimal graft quality

In summary, this study uncovered new insights into graft function related proteins in liver allografts. The L-FABP level might be an effective biomarker for evaluating donor quality in liver transplantation.

## Abbreviations

ALT: alanine transaminase;
AST: aspartate aminotransferase;
ALP: alkaline phosphatase;
BMI: body mass index;
COG: clusters of orthologous groups of proteins;
DCD: donation after cardiac death;
EAD: early allograft dysfunction;
ECD: extended criteria donors;
GO: gene ontology;
HOPE: hypothermic oxygenated machine perfusion;
iTRAQ: isobaric Tags for Relative and Absolute Quantification;
IRI: ischemia reperfusion injury;
KEGG: kyoto encyclopedia of genes and genomes;
L-FABP: liver fatty acid binding protein;
LDH: lactate degyfrogenase;
LT: liver transplantation;
MRM: multiple reaction monitoring;
NMP: normothermic machine perfusion;
OG: optimal graft;
PNF: primary nonfunction;
SDS-PAGE: sodium dodecyl sulfate – polyacrylamide gel electrophoresis;
TBIL: total bilirubin;

## Acknowledgements

The authors wish to acknowledge the assistance of BGI-Shenzhen for the study monitoring, analysis and data interpretation. This work was supported by the National Natural Science Foundation of China (81401324 and 81770410), Guangdong Basic and Applied Basic Research Foundation (2020A1515011557), Science and Technology Planning Project of Guangdong Province (2016A020215048), Guangdong Provincial Key Laboratory of Organ Donation and Transplant Immunology (2013A061401007), Guangdong Provincial International Cooperation Base of Science and Technology (Organ Transplantation) (2015B050501002), Scientific Program for Young teacher of Sun Yat-sen University (16ykpy05), China.

## References

1. Lee, D. D., Singh, A., Burns, J. M., Perry, D. K., Nguyen, J. H., and Taner, C. B. (2014) Early Allograft Dysfunction in Liver Transplantation With Donation After Cardiac Death Donors Results in lnferior Survival. Liver Transplant 20, 1447–1453

2. De Carlis, R., Di Sandro, S., Lauterio, A., Botta, F., Ferla, F., Andorno, E., Bagnardi, V., and De Carlis, L. (2018) Liver Grafts From Donors After Circulatory Death on Regional Perfusion With Extended Warm lschemia Compared With Donors After Brain Death. Liver Transplant 24, 1523–1535

3. Olthoff, K. M., Kulik, L., Samstein, B., Kaminski, M., Abecassis, M., Emond, J., Shaked, A., and Christie, J. D. (2010) Validation of a current definition of early allograft dysfunction in liver transplant recipients and analysis of risk factors. Liver Transpl 16, 943–949

4. Pareja, E., Cortes, M., Hervas, D., Mir, J., Valdivieso, A., Castell, J. V., and Lahoz, A. (2015) A score model for the continuous grading of early allograft dysfunction severity. Liver Transpl 21, 38–46

5. Corradini, S. G., Elisei, W., De Marco, R., Siciliano, M., lappelli, M., Pugliese, F., Ruberto, F., Nudo, F., Pretagostini, R., Bussotti, A., Mennini, G., Eramo, A., Liguori, F., Merli, M., Attili, A. F., Muda, A. O., Natalizi, S., Berloco, P., and Rossi, M. (2005) Preharvest donor hyperoxia predicts good early graft function and longer graft survival after liver transplantation. Liver Transpl 11, 140–151

6. Zhao, Q., Guo, Z., Deng, W., Fu, S., Zhang, C., Chen, M., Ju, W., Wang, D., and He, X. (2016) Calpain 2-mediated autophagy defect increases susceptibility of fatty livers to ischemia-reperfusion injury. Cell Death Dis 7, e2186

7. Suzuki, S., Toledo-Pereyra, L. H., Rodriguez, F. J., and Cejalvo, D. (1993) Neutrophil infiltration as an important factor in liver ischemia and reperfusion injury. Modulating effects of FK506 and cyclosporine. Transplantation 55, 1265–1272

8. Nemes, B., Gaman, G., Polak, W. G., Gelley, F., Hara, T., Ono, S., Baimakhanov, Z., Piros, L., and Eguchi, S. (2016) Extended-criteria donors in liver transplantation Part ll: reviewing the impact of extended-criteria donors on the complications and outcomes of liver transplantation. Expert Rev Gastroenterol Hepatol 10, 841–859

9. Marcon, F., Schlegel, A., Bartlett, D. C., Kalisvaart, M., Bishop, D., Mergental, H., Roberts, K. J., Mirza, D. F., lsaac, J., Muiesan, P., and Perera, M. T. (2018) Utilization of Declined Liver Grafts Yields Comparable Transplant Outcomes and Previous Decline Should Not Be a Deterrent to Graft Use. Transplantation 102, E211–E218

10. Bacchella, T., Galvao, F. H., Jesus de Almeida, J. L., Figueira, E. R., de Moraes, A., and Cesar Machado, M. C. (2008) Marginal grafts increase early mortality in liver transplantation. Sao Paulo Med J 126, 161–165

11. Czigany, Z., Schoning, W., Ulmer, T. F., Bednarsch, J., Amygdalos, l., Cramer, T., Rogiers, X., Popescu, l., Botea, F., Fronek, J., Kroy, D., Koch, A., Tacke, F., Trautwein, C., Tolba, R. H., Hein, M., Koek, G. H., Dejong, C. H. C., Neumann, U. P., and Lurje, G. (2017) Hypothermic oxygenated machine perfusion (HOPE) for orthotopic liver transplantation of human liver allografts from extended criteria donors (ECD) in donation after brain death (DBD): a prospective multicentre randomised controlled trial (HOPE ECD-DBD). BMJ Open 7, e017558

12. Ghinolfi, D., Rreka, E., De Tata, V., Franzini, M., Pezzati, D., Fierabracci, V., Masini, M., Cacciatoinsilla, A., Bindi, M. L., Marselli, L., Mazzotti, V., Morganti, R., Marchetti, P., Biancofiore, G., Campani, D., Paolicchi, A., and De Simone, P. (2019) Pilot, Open, Randomized, Prospective Trial for Normothermic Machine Perfusion Evaluation in Liver Transplantation From Older Donors. Liver Transpl 25, 436–449

13. Pavel, M. C., Reyner, E., Molina, V., Garcia, R., Ruiz, A., Roque, R., Diaz, A., Fuster, J., and Garcia-Valdecasas, J. C. (2019) Evolution Under Normothermic Machine Perfusion of Type 2 Donation After Cardiac Death Livers Discarded as Nontransplantable. J Surg Res 235, 383–394

14. Xie, H., Zhang, L., Guo, D., Yang, Z., Zhu, H., Zhou, K., Feng, X., Wei, Q., Xu, X., Song, P., Wen, X., Li, J., Liu, J., and Zheng, S. (2019) Protein Profiles of Pretransplant Grafts Predict Early Allograft Dysfunction After Liver Transplantation From Donation After Circulatory Death. Transplantation

15. Feng, Q., Zhao, N., Xia, W., Liang, C., Dai, G., Yang, J., Sun, J., Liu, L., Luo, L., and Yang, J. (2019) lntegrative proteomics and immunochemistry analysis of the factors in the necrosis and repair in acetaminophen-induced acute liver injury in mice. J Cell Physiol 234, 6561–6581

16. Pelsers, M. M., Morovat, A., Alexander, G. J., Hermens, W. T., Trull, A. K., and Glatz, J. F. (2002) Liver fatty acid-binding protein as a sensitive serum marker of acute hepatocellular damage in liver transplant recipients. Clin Chem 48, 2055–2057

17. Tajima, S., Yamamoto, N., and Masuda, S. (2019) Clinical prospects of biomarkers for the early detection and/or prediction of organ injury associated with pharmacotherapy. Biochem Pharmacol 170, 113664

18. Wang, G., Gong, Y., Anderson, J., Sun, D., Minuk, G., Roberts, M. S., and Burczynski, F. J. (2005) Antioxidative function of L-FABP in L-FABP stably transfected Chang liver cells. Hepatology 42, 871–879

19. Gyamfi, M. A., He, L., French, S. W., Damjanov, l., and Wan, Y. J. (2008) Hepatocyte retinoid X receptor alpha-dependent regulation of lipid homeostasis and inflammatory cytokine expression contributes to alcohol-induced liver injury. J Pharmacol Exp Ther 324, 443–453

20. Ek-Von Mentzer, B. A., Zhang, F., and Hamilton, J. A. (2001) Binding of 13-HODE and 15-HETE to phospholipid bilayers, albumin, and intracellular fatty acid binding proteins. implications for transmembrane and intracellular transport and for protection from lipid peroxidation. J Biol Chem 276, 15575–15580

21. Divine, J. K., McCaul, S. P., and Simon, T. C. (2003) HNF-1alpha and endodermal transcription factors cooperatively activate Fabpl: MODY3 mutations abrogate cooperativity. Am J Physiol Gastrointest Liver Physiol 285, G62–72

22. Newberry, E. P., Xie, Y., Lodeiro, C., Solis, R., Moritz, W., Kennedy, S., Barron, L., Onufer, E., Alpini, G., Zhou, T., Blaner, W. S., Chen, A., and Davidson, N. O. (2019) Hepatocyte and stellate cell deletion of liver fatty acid binding protein reveals distinct roles in fibrogenic injury. FASEB J 33, 4610–4625

23. Pelsers, M. M., Namiot, Z., Kisielewski, W., Namiot, A., Januszkiewicz, M., Hermens, W. T., and Glatz, J. F. (2003) lntestinal-type and liver-type fatty acid-binding protein in the intestine. Tissue distribution and clinical utility. Clin Biochem 36, 529–535

24. Mikus, M., Drobin, K., Gry, M., Bachmann, J., Lindberg, J., Yimer, G., Aklillu, E., Makonnen, E., Aderaye, G., Roach, J., Fier, l., Kampf, C., Gopfert, J., Perazzo, H., Poynard, T., Stephens, C., Andrade, R. J., Lucena, M. l., Arber, N., Uhlen, M., Watkins, P. B., Schwenk, J. M., Nilsson, P., and Schuppe-Koistinen, l. (2017) Elevated levels of circulating CDH5 and FABP1 in association with human drug-induced liver injury. Liver lnt 37, 132–140

25. Akbal, E., Koklu, S., Kocak, E., Cakal, B., Gunes, F., Basar, O., Tuna, Y., and Senes, M. (2013) Liver fatty acid-binding protein is a diagnostic marker to detect liver injury due to chronic hepatitis C infection. Arch Med Res 44, 34–38

26. van den Broek, M. A. J., Bloemen, J. G., Dello, S. A. W. G., van de Poll, M. C. G., Damink, S. W. M. O., and Dejong, C. H. C. (2011) Randomized controlled trial analyzing the effect of 15 or 30 min intermittent Pringle maneuver on hepatocellular damage during liver surgery. Journal of Hepatology 55, 337–345

27. van de Poll, M. C., Derikx, J. P., Buurman, W. A., Peters, W. H., Roelofs, H. M., Wigmore, S. J., and Dejong, C. H. (2007) Liver manipulation causes hepatocyte injury and precedes systemic inflammation in patients undergoing liver resection. World J Surg 31, 2033–2038

